# Nanoparticle-mediated transgene expression of *insulin-like growth factor 1* in the growth restricted guinea pig placenta increases placenta nutrient transporter expression and fetal glucose concentrations

**DOI:** 10.1101/2021.06.24.449769

**Authors:** Rebecca L. Wilson, Kristin Lampe, Mukesh K. Gupta, Craig L. Duvall, Helen N. Jones

## Abstract

Fetal growth restriction (FGR) significantly contributes to neonatal and perinatal morbidity and mortality. Currently, there are no effective treatment options for FGR during pregnancy. We have developed a nanoparticle gene therapy targeting the placenta to increase expression of human *insulin-like growth factor 1* (*hIGF-1*) to correct fetal growth trajectories. Using the maternal nutrient restriction (MNR) guinea pig model of FGR, an ultrasound-guided, intra-placental injection of non-viral, polymer-based nanoparticle gene therapy containing plasmid with the *hIGF-1* gene and placenta-specific *Cyp19a1* promotor was administered at mid-pregnancy. Sustained *hIGF-1* expression was confirmed in the placenta five days after treatment. Whilst gene therapy treatment did not change fetal weight, circulating fetal glucose concentration were 33-67% higher. This was associated with increased expression of glucose and amino acid transporters in the placenta. Additionally, nanoparticle gene therapy treatment increased the fetal capillary volume density in the placenta, and reduced interhaemal distance between maternal and fetal circulation. Overall, our findings, that gene therapy treatment results in changes to glucose transporter expression and increases fetal glucose concentrations within a short time period, highlights the translational potential this treatment could have in correcting impaired placental nutrient transport in human pregnancies complicated by FGR.

## Introduction

Fetal growth restriction (FGR: estimated fetal weight <10^th^ percentile) occurs in up to 10% of pregnancies with suboptimal fetal nutrition and uteroplacental perfusion accounting for 25-30% of cases [1, 2]. Perinatal outcomes depend largely on the severity of FGR, however, and FGR is a leading cause of infant morbidity and mortality [3]. In addition, FGR has been implicated in contributing to the development of long-term health outcomes including increasing the risk for future cardiovascular and endocrine diseases such as diabetes [4] as well as being associated with cognitive and learning disabilities [5]. Currently, there is limited preventative strategies and no effective treatment options. However, advances in nanomedicine are allowing for the development of potential therapeutics to treat the placenta and correct fetal growth trajectories [6].

Currently, there are a number of studies being undertaken that are designed at specifically targeting the placenta to improve pregnancy outcome [7, 8]. Furthermore, advances in nanoparticle capabilities, which exploit existing cellular uptake mechanisms, are allowing for the delivery of drugs, gene therapies and other pharmacological agents with maximized efficiency in smaller doses [9, 10]. We are developing the use of a polymer-based nanoparticle gene therapy that facilitates non-viral gene delivery specifically to the placenta [11, 12]. Using a synthetic HPMA-DMEAMA (N-(2-hydroxypropyl) methacrylamide-2-(dimethylamino)ethyl methacrylate) diblock copolymer, complexed with a plasmid containing the human *insulin-like 1 growth factor* (*hIGF-1*) gene under the control of placenta trophoblast specific promotors (*PLAC1* and *CYP19A1*), we have shown efficient nanoparticle uptake into human syncytiotrophoblast ex vivo [12] and increased placenta expression of *hIGF-1* that is capable of maintaining fetal growth in a surgically-induced mouse model of FGR [11]. Most importantly, we have shown this technology to be safe to both mother and fetus, as both the nanoparticle and plasmid do not cross the placental syncytium, and treatment has never been associated with maternal morbidities, or increased fetal loss [11-13].

Both active and passive nutrient transport systems maintain placental nutrient supply to the fetus [14]. In humans, maternal-fetal exchange of oxygen, nutrients and waste occurs across the syncytiotrophoblasts (the multi-nucleated syncytium that serves as the primary barrier between maternal and fetal blood) and fetal capillary endothelium [15]. Whilst the fetal capillary endothelium largely allows unrestricted transfer of small molecules and ions, it is the syncytiotrophoblasts which express the array of nutrient transporter proteins critical of mediating active transport of nutrients and waste [16]. In pregnancies complicated by FGR, reduced placental glucose and amino acid transport capacities has consistently been shown, and animal models report downregulation of placenta transport systems prior to the development of FGR [15]. Changes in nutrient transporter capabilities in the placenta are often associated with dysregulation of metabolic hormones, like IGF-1, which co-ordinate the flux of glucose, amino acids and lipids. IGF-1 is a secreted protein produced by the placenta throughout pregnancy [17]. Through binding with the IGF-1 Receptor (IGF-1R), it regulates numerous aspects of placenta function including metabolism, mitogenesis, and differentiation [18]. Furthermore, we and others have demonstrated that increased placental expression of IGF-1 is capable of upregulating glucose and amino acid transport to the fetus and thus modulating fetal growth [19-21].

Development of a treatment for FGR is made more difficult by the lack of definitive ways to predict FGR prior to diagnosis, and as such, effective therapies must be capable of correcting the effects of existing placental dysfunction. Guinea pigs offer a unique advantage in studying the placenta, fetal development, and reproductive health as they have similar developmental milestones to humans both throughout gestation and following birth [22, 23]. Guinea pig placentas better reflect humans over other small animal species [24], exhibit deeper placental trophoblast invasion [25], and undergo changes to the maternal hormonal profile throughout pregnancy that more closely resemble humans [26]. Additionally, non-invasive, maternal nutrient restriction (MNR) where dam food intake is restricted to 70% 4 weeks prior to pregnancy until mid-gestation and 90% thereafter, induces FGR [27, 28]. Here, we aimed to characterize placenta uptake of nanoparticle and the effect of gene therapy on placenta morphology and glucose transport in the guinea pig MNR model of FGR.

## Materials and Methods

### Synthesis of poly(2-hydroxypropyl methacrylamide (PHPMA_115_-ECT))

In a 25 mL round bottom flask equipped with a magnetic stirrer bar, ECT (4-cyano-4-(ethylsulfanylthiocarbonyl) sulfanylvpentanoic acid; 0.025 mmol, 0.006 g), HPMA (5 mmol, 0.715 g), and AIBN (2,2′-azo-*bis*-isobutyrylnitrile; 0.0025 mmol, 0.0004 g) were dissolved in DMF (dimethyl formamide; 5 mL) and sealed with septa. The solution was degassed for 30 mins by purging with ultra-high purity nitrogen and submerged into a preheated oil bath at 65 °C. After 24 h, the crude polymerization mixture was purified through precipitation into cold diethyl ether. The pale-yellow color polymer was dried overnight under vacuum and the structure of the polymer was characterized by ^1^H NMR spectroscopy.

### Synthesis of diblock copolymer of poly(2-hydroxypropyl methacrylamide-*b*-N, N-Dimethyl aminoethyl methacrylate) (PHPMA_115_-*b*-PDMAEMA_115_)

RAFT (reversible addition fragmentation chain transfer) polymerization of DMEAMA was performed using PHPMA_115_-ECT as a macro-CTA (chain transfer agent) and AIBN as a free radical initiator. A solution of PHPMA_115_-ECT (0.031 mmol, 0.51 g), DMAEMA (4.65 mmol, 0.73 g, 0.782 mL), and AIBN (0.0031 mmol, 0.0005 g) in methanol (5 mL) was degassed for 30 minutes through purging with high purity nitrogen. To initiate the polymerization, the reaction flask was submerged in preheated oil at 70 °C. The crude polymerization mixture was precipitated three times into cold pentane and dried under high vacuum after 24 h. The precipitated polymer was dissolved into methanol and dialyzed first against methanol for 24 h and then against deionized water for another 48 h. The dialyzed polymer was lyophilized to obtain a white powder, and formation of the diblock copolymer was confirmed by ^1^H NMR spectroscopy.

### Nanoparticle Formation

Nanoparticles were formed by complexing plasmids containing either the *GFP* or *hIGF-1* gene under control of placental trophoblast specific promotors *PLAC1* or *CYP19A1* with a non-viral PHPMA_115_-b-PDMEAMA_115_ co-polymer. All plasmids used were cloned from a pEGFP-C1 plasmid (Clonetech Laboratories, Mountain View, CA), replacing the *CMV* promotor with either the *PLAC1* or *CYP19A1* promotor, and the *GFP* gene with *hIGF1*. The dialyzed polymer was resuspended in sterile water at a concentration of 10 mg/mL, then mixed with plasmid at a ratio of 1.5 µL polymer to 1 µg plasmid, and made to a total injection volume of 200 µL with sterile phosphate buffered saline (PBS). Characterization of the physiochemical properties, and cellular safety and efficiency of the PHPMA_115_-b-PDMEAMA_115_ co-polymer complexed with the pEGFP-C1 plasmid to form the nanoparticle has been published previously [29]. To initially determine nanoparticle uptake and promotor recognition in the guinea pig placenta, polymer was conjugated with a Texas Red fluorophore, described previously [12].

### Animal Care and Transuterine, Intra-placental Nanoparticle Injections

Female (dams) Dunkin-Hartley guinea pigs were purchased (Charles River Laboratories, Wilmington, MA) at 500-550 g and housed individually in a temperature-controlled facility with a 12 h light-dark cycle. Whilst acclimatizing to the facilities and for initial investigations into nanoparticle uptake in the placenta, food (LabDiet diet 5025: 27% protein, 13.5% fat, and 60% carbohydrate as % of energy) and water was provided ad libitum. Animal care and usage was approved by the Institutional Animal Care and Use Committees at Cincinnati Children’s Hospital and Medical Center (Protocol number 2017-0065) and University of Florida (Protocol number 202011236). See schematic S1 in Supplemental Material for outline of experimental design.

In the initial studies to determine nanoparticle uptake and transgene expression in the guinea pig placenta, the Texas Red conjugated polymer was complexed with plasmid containing the *GFP* gene. Dams (*n* = 6; 2 sham, 2 *Plac1* promotor, 2 *Cyp19a1* promotor) were time mated through daily monitoring of the estrus cycle by inspecting perforation of the vaginal membrane [30]. Ovulation was assumed to occur when the vaginal membrane was fully perforated and designated gestational day (GD) 1. Pregnancy confirmation ultrasounds were performed at GD21. At GD30 (mid-pregnancy), dams underwent an ultrasound-guided, transuterine, intra-placental nanoparticle injection (containing 60 µg plasmid in 200 µL injection) or sham injection (200 µL of PBS) using a 32G x 1-inch needle placed into the placental labyrinth as close to the visceral cavity as possible. This time point was chosen because this is the earliest point in the guinea pig pregnancy where fetuses are growth restricted. Dams were anaesthetized using isoflurane, and their abdomen shaved and cleaned using isopropanol. Ultrasounds were performed using a Voluson I portable ultrasound machine (GE) with a 125 E 12 MHz vascular probe (GE). The abdominal cavity was scanned to determine the number and location of fetuses within the uterine horns. Where possible, the uterine horn which contained only a single fetus was chosen to receive the nanoparticle treatment. The fetuses on the opposite horn were left untreated, keeping consistent with our previous mouse studies [11, 13]. On the minimal occasion that all fetuses were located in the same uterine horn, the placenta of the fetus located closest to the cervix was injected. Following injection, the placenta was briefly monitored using the ultrasound for signs of hemorrhage and fetal heart beat monitored to ensure no immediate fetal demise. Dams were then allowed to recover in a humidicrib maintained at 37ºC before being return to their home cages. Euthanasia occurred 30 hours after injection by carbon dioxide asphyxiation followed by cardiac puncture and exsanguination. The gravid uterus was dissected and fetuses and maternal-fetal interface weighed. Maternal and fetal organs including heart, lungs, liver, kidney and brain were collected. These tissues, along with the maternal-fetal interface, were halved and either fixed in 4% w/v paraformaldehyde (PFA) or snap-frozen in liquid nitrogen and stored at -80ºC. Tissues placed in PFA were left at 4ºC until appropriately fixed, transferred to 30% w/v sucrose solution and then rapidly frozen in OCT and stored at -80ºC.

To determine the effect of gene therapy treatment on placental function under adverse maternal conditions, dams were weighed, ranked from heaviest to lightest, and systematically assigned to either the control diet group (n = 7) where food and water was provided ad libitum, or maternal nutrient restricted group (MNR; n = 12) so that mean weight was no different between the control and MNR groups. MNR dams were provided water ad libitum however, food intake was restricted to 70% per kilogram body weight of the control group from at least four weeks preconception through to mid-pregnancy (GD30), thereafter increasing to 90% [27, 28]. Time mating and pregnancy confirmation ultrasounds were performed as previously described; dams not confirmed pregnant were re-mated with all but one dam falling pregnant within three mating attempts. At GD30-33, MNR dams underwent an ultrasound-guided, transuterine, intra-placental nanoparticle injection of plasmid containing *hIGF-1*, or sham injection (n = 7 control and 5 MNR) as previously described. Inaccessibility to surgical facilities over weekends prevented injections being performed at the exact same time point in pregnancy. Plasmid expression of *hIGF1* is transient as the plasmid does not integrate into the genome. We have shown in mice placentas *in vivo*, a decline in the presence of nanoparticle, based on Texas-red fluorescence, with the placenta [31] but robust mRNA expression of plasmid-specific *hIGF1* at 96 h after injection [11]. Therefore, we chose to sacrifice all dams five days after injection (GD35-38) to ensure that transient expression of plasmid *hIGF1* was still present within the placenta. On the day of sacrifice, dams were fasted for four hours, weighed and then euthanized as previously described. Glucose concentrations in both maternal and fetal blood was measured using a glucometer. Tissue collection was performed as previously described however, tissue placed in PFA was processed and paraffin embedded following standard protocols. Fetal sex was determined by examination of the gonads and confirmed using PCR as previously published [32].

### Fluorescent Microscopy of Nanoparticle Uptake

Placenta and fetal liver tissue from dams treated with the Texas Red-*GFP* nanoparticle were assessed microscopically for nanoparticle uptake and transgene expression. 7 µm thick tissue sections were obtained, OCT was cleared by washing in ice-cold PBS, nuclei counterstained with DAPI (*Invitrogen*), and mounted using Prolong Gold Antifade mounting medium (*Invitrogen*). Texas Red and GFP fluorescence was visualized using the Nikon Eclipse Ti Inverted microscope.

Exposure time was set using sham treated tissue to eliminate auto- and background fluorescence and thus, any fluorescence observed was due to the presence of nanoparticle (indicated by Texas-Red fluorescence) or plasmid expression (indicated by GFP fluorescence).

### In situ Hybridization for Plasmid-Specific mRNA

In situ hybridization of plasmid-specific mRNA was performed on maternal-fetal interface tissue treated with nanoparticle gene therapy to confirm sustained transgene expression five days after injection as previously described [12, 32]. Two 5 µm thick, paraffin embedded serial tissue sections were obtained. For the first section, in situ hybridization was performed using a BaseScope™ probe (*Advanced Cell Diagnostics*) custom designed to recognize the sequence between the stop codon and polyA signaling of the plasmid, following standard protocol. A section of human placenta explant treated with nanoparticle was included as a positive control. For the second section, immunohistochemistry (IHC) was performed to confirm localization of plasmid-specific mRNA in placental trophoblast cells following standard IHC protocols. Following de-waxing and rehydrating, targeted antigen retrieval was performed using boiling 1x Targeted Retrieval Solution (*Dako*) for 20 mins. Endogenous peroxidase activity was suppressed by incubating sections in 3% hydrogen peroxide for 10 mins and then sections were blocked in 10% goat serum with 1% bovine serum albumin (BSA; *Jackson ImmunoResearch*) in PBS for 1 h at room temperature. Anti-pan cytokeratin (*Sigma C2562*; 1:200) primary antibody was applied overnight at 4ºC followed by an incubation with a biotinylated anti-mouse IgG secondary antibody (*Vector BA-9200*; 1:200) for 1 h at room temperature. The Vector ABC kit (*Vector*) was used to amplify staining and detected using DAB (*Vector*) for brown precipitate. Hematoxylin was used to counterstain nuclei. In situ hybridization was also performed on fetal liver sections to confirm the inability for plasmid to cross the placenta and enter fetal circulation five days after treatment. All sections were imaged using the Axioscan scanning microscope (*Zeiss*).

### Placenta Morphological Analysis

For placenta morphometric analyses, 5 µm thick, full-face sections of maternal-fetal interface were stained with Masson’s trichrome stain and double-label immunohistochemistry (IHC). Sections were de-waxed and rehydrated following standard protocols. To visualize the placenta, sub-placenta and decidual regions, Masson’s trichrome (*Sigma*) was performed as per manufacturers specifications. Whole section imaging was performed using the Axioscan Scanning Microscope (*Zeiss*) and placenta and sub-placenta/decidua areas were measured using the Zen Imaging software (*Zeiss*). For double-label IHC, fetal capillary endothelium and placental trophoblasts were distinguished using anti-vimentin (*Dako Vim3B4*: 1:100) and anti-pan cytokeratin (*Sigma C2562*; 1:200) antibodies, respectively [33] (Supplemental Fig. S1). Antigen retrieval was performed with 0.03% protease (*Sigma*) for 15 min at 37ºC and endogenous peroxide activity was blocked with 3% hydrogen peroxide for 30 mins. The sections were then blocked with serum-free protein block (*Dako*) for 10 mins before anti-vimentin antibody was applied to slides, diluted in 10% guinea pig serum and 1% BSA, and left overnight at 4ºC. Biotinylated anti-mouse IgG secondary antibody (*Vector BA-9200*; 1:200) was diluted in 10% guinea pig serum with 1% BSA and applied to sections for 30 mins. Antibody staining was amplified using the Vector ABC kit (*Vector*) for 40 mins and visualized using 3, 3′ diaminobenzidine tetrahydrochloride (DAB; *Vector*) with nickel to create a black precipitate. This process, from the protein block, was then repeated for anti-pan cytokeratin, however the anti-pan cytokeratin antibody remained on the sections for 2 h at room temperature and nickel was omitted from the DAB so as to form a brown precipitate. Counterstaining occurred using hematoxylin. For detailed analysis of the labyrinthine structure, the proportion of trophoblast, fetal capillaries and maternal blood space was calculated using point counting with an isotropic L-36 Merz grid [34]. Double-labeled sections were imaged using the Axioscan (*Zeiss*) and then 10 random 40x fields of view were captured. Each field of view was then counted as described in [34]. Interhaemal distance of the maternal blood space and between the maternal blood space and fetal capillaries was also calculated using the Merz grid and line intercept counting. Within-assay variation of <5% was confirmed with repeated measurement.

### RNA Isolations and Quantitative PCR (qPCR)

For gene expression analysis, approximately 150-200 mg of snap frozen placenta tissue was lysed in RLT-lysis buffer (*Qiagen*) with homogenization aided by a tissue-lyzer. RNA was extracted using the RNeasy Midi kit (*Qiagen*), and included DNase treatment following standard manufacturers protocol. 1 µg of RNA was converted to cDNA using the High-capacity cDNA Reverse Transcription kit (*Applied Biosystems*) and diluted to 1:100. For qPCR, 2.5 µL of cDNA was mixed with 10 µL of PowerUp SYBR green (*Applied Biosystems*), 1.2 µL of primers (See supplemental table S1 for primer sequences) at a concentration of 10 nM, and water to make up a total reaction volume of 20 µL. Gene expression was normalized using housekeeping genes *β-actin* and *Rsp20*. Reactions were performed using the StepOne-Plus Real-Time PCR System (*Applied Biosystems*), and relative mRNA expression calculated using the comparative CT method with the StepOne Software v2.3 (*Applied Biosystems*).

### Placenta Western Blot

For analysis of glucose transporter protein expression, placenta tissue was homogenized in ice-cold RIPA buffer containing protease and phosphatase inhibitors and protein concentrations determined using Pierce™ Coomassie Plus Assay Kit (*Thermo Scientific*) following standard protocol. 40 µg of protein was then mixed with Nupage SDS Loading Buffer (*Invitrogen*) and Reducing Agent (*Invitrogen*) and denatured by heating at 95ºC for 10 min. The lysates were then run on a 10% Tris-Bis precast gel (*Invitrogen*) following manufacturers protocols and transferred onto nitrocellulose membranes using the Bolt Mini Electrophoresis unit (*Invitrogen*). A Prestain Protein ladder (PageRuler, *Thermo Scientific*) was included on the gels. Membranes were placed into 5% skim-milk in Tris-buffered Saline containing Tween 20 (TBS-T) and incubated overnight at 4ºC. Primary antibodies (Slc2A1: *Abcam ab15309* 1:500, Slc2A3: *Abcam ab15311* 1:200, Slc38A1: *Abcam ab104684* 1:200, Slc38a2: *Abcam ab90677* 1:200) were then applied for 1-2 h at room temperature, the membranes washed 3 times in fresh TBS-T, and then further incubated with a HRP conjugated secondary (*Cell Signaling* 1:1000) for 1 h at room temperature. Protein bands were visualized by chemiluminescence using SuperSignal West Femto Maximum Sensitivity Substrate (*Thermo Scientific*) on the Universal Imager (*Biorad*). Signal intensity of the protein bands were calculated using Image Studio Lite (version 5.2, *Licor*) and normalized to β-actin.

Localization of Slc2A1 in the syncytiotrophoblast membranes was visualized using IHC as described in *In situ Hybridization for Plasmid-Specific mRNA* section. Primary antibody for Slc2A1 (*Abcam ab15309*) was diluted 1:750 and applied to sections overnight at 4ºC, and biotinylated anti-rabbit IgG secondary antibody (*Vector BA-1000*; 1:200) also used as previously described.

### Statistical Analysis

All statistical analyses were performed using SPSS Statistics 26 software. Generalized estimating equations were used to determine differences between diet and nanoparticle treatment. Dams were considered the subject, diet, nanoparticle treatment and fetal sex treated as main effects, maternal environment treated as a random effect and gestational age as a covariate. Litter size was also included as a covariate but removed as there was no significant effect for any of the outcomes. Statistical significance was considered at P≤0.05. For statistically significant results, a Bonferroni post hoc analysis was performed. Results are reported as estimated marginal means ± standard error.

## Results

### Synthesis and characterization of PHPMA_115_-b-PDMAEMA_115_

The general schematic for the synthesis of the diblock copolymer is outlined in scheme S2 (Supplemental Material). The molar ratio of chain transfer agent (CTA) and 2,2′-azo-*bis*-isobutyrylnitrile (AIBN) was 10:1. ^1^H NMR spectra showed the presence of all characteristic peaks for PHPMA_115_-ECT (Supplemental Fig. S2). The PHPMA macro-CTA was then chain extended by RAFT (reversible addition fragmentation chain transfer) block polymerization of PDMAEMA to yield the final diblock copolymer PHPMA_115_-b-PDMAEMA_115_. The presence of all characteristic peaks for PHPMA and PDMAEMA in ^1^H NMR spectra confirmed successful diblock copolymer formation (Supplemental Fig. S2).

### The nanoparticle is rapidly endocytosed in the guinea pig placenta but only the *Cyp19a1* promotor is recognized

In order to confirm nanoparticle uptake and promotor recognition in the guinea pig placenta, a plasmid containing the green fluorescent protein (*GFP*) gene was complexed with a Texas Red conjugated HPMA-DMAEMA co-polymer. Texas Red fluorescence was observed in the sub-placental/decidual region of the guinea pig placenta 30 h after treatment indicating the ability of nanoparticle to enter placental cells (Supplemental Fig. S3). However, GFP fluorescence was only observed in placental cells of tissue that received an inject of nanoparticle containing a plasmid with the *Cyp19a1* promotor and not the *Plac1* promotor (Supplemental Fig. S3). GFP fluorescence appeared to be localized in vesicles within the cytoplasm of cells. There was no Texas-red fluorescence observed in the fetal liver confirming the inability for the nanoparticle to enter fetal circulation. Using qPCR, we have previously shown in mice placentas, expression of *hIGF-1* following nanoparticle treatment [11, 12]. Given the high degree of homology between human and guinea pig *Igf-1*, human-specific, and therefore nanoparticle gene therapy specific, *hIGF-1* expression could not be determined. However, *in situ* hybridization analysis of placenta tissue receiving the *Cyp19a1-hIGF-1* nanoparticle five days after intra-placental injection, confirmed the presence of plasmid-specific mRNA in the trophoblasts of the sub-placenta/decidua region (Fig. 1A). *In situ* hybridization in sham injected placentas showed no positive plasmid-specific mRNA expression, as expected (Fig 1B). Unlike in the mouse studies, in which untreated placentas in the opposite uterine horn to treated placentas did not show increased *hIGF-1* expression due to nanoparticle gene therapy treatment [11, 13], plasmid-specific mRNA was not just confined to the placenta that received the intra-placental injection, but was found in all placentas of the litter (Supplemental Fig. S4). No positive ISH staining was found in the fetal liver further confirming the inability for plasmid to cross the placental barrier and enter fetal circulation after five days (Supplemental Fig. S5). Analysis of guinea pig *Igf-1* mRNA revealed no expression (amplification at CT>37) in the guinea pig placenta labyrinth whilst sub-placenta/decidua expression of *Igf-1* was not altered by either diet or nanoparticle treatment (P>0.05; Supplemental Fig. S5A). Igf-1 protein, which is secreted and therefore not expected to be in the cytoplasm of cells, could not be detected in placenta tissue using IHC or Western Blot analyses (Supplemental Fig. S6B-C).

**Fig. 1.**
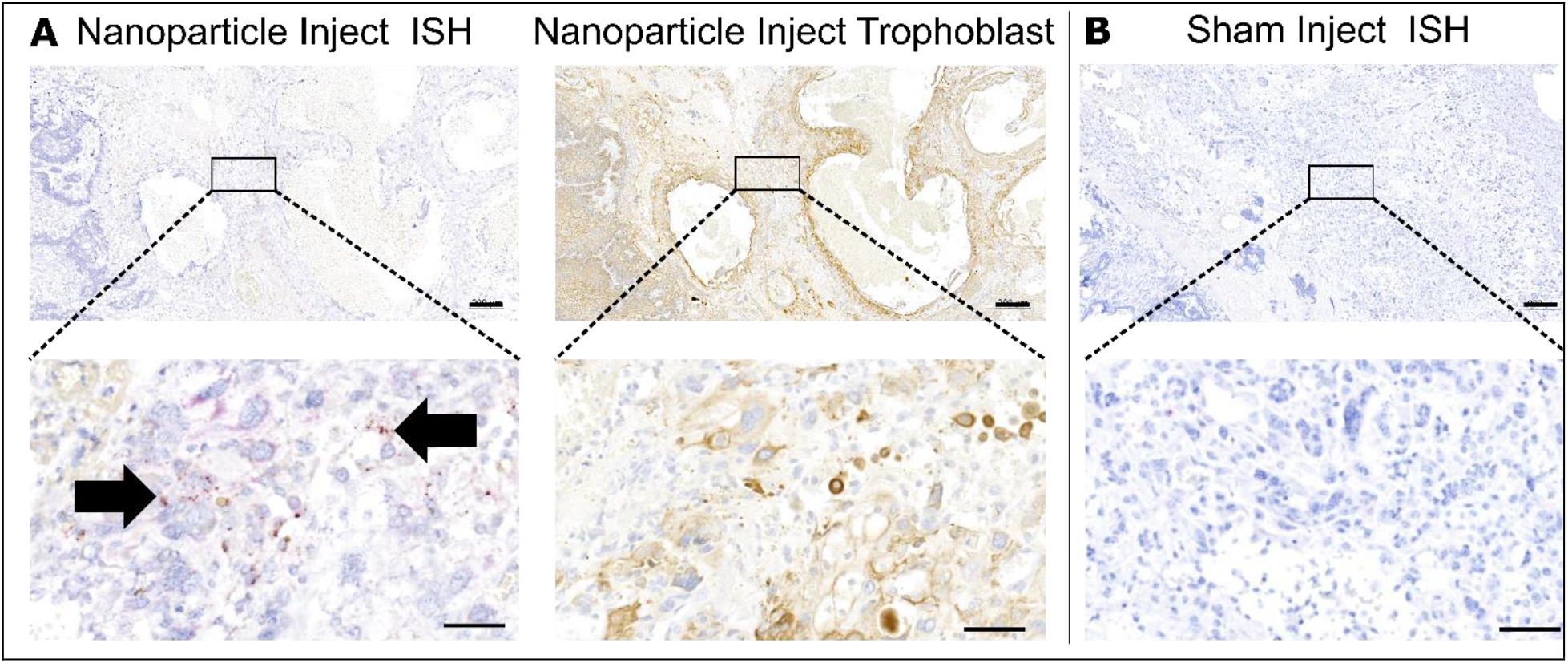
In situ hybridization (ISH) for plasmid-specific mRNA expression in the guinea pig placenta five days after intra-placental nanoparticle treatment. **A**. ISH (first panel) confirmed plasmid-specific mRNA expression in the guinea pig sub-placenta/decidua five days after nanoparticle treatment. Serial sectioning and immunohistochemistry (second panel) confirmed plasmid-specific mRNA was localized to trophoblast cells. **B**. Sham injected tissue was used as a negative control and no positive staining for plasmid-specific mRNA was observed. Representative images at low magnification (top row) and high magnification (bottom row). Arrows = the presence of red dots indicating plasmid-specific mRNA. Scale bar top row = 200 µm, scale bar bottom row = 50 µm

### Short-term placenta nanoparticle treatment was not toxic to fetal development

Only 1 dam (MNR, 3 viable fetuses and 1 resorption) was recorded as having a resorption. The resorption appeared as an amorphous, dark lump within the uterus indicative of early pregnancy loss not associated with nanoparticle gene therapy treatment. Furthermore, there was no difference in litter size between the MNR dams compared to the control *ad libitum* fed dams (mean ± SEM: control = 3.2 ± 0.35 vs. restricted = 2.8 ± 0.24, P>0.05). Maternal carcass weight (maternal weight minus fetal and maternal-fetal interface weight) was lower in MNR dams compared to control (P=0.03; Fig. 2A), but not different between MNR-sham and MNR-nanoparticle gene therapy treated (P>0.05). There was also no difference in maternal liver and spleen weight as a percentage of carcass weight with either diet or nanoparticle treatment (Fig. 2B and 2C, respectively). Fetal weight was approximately 22-25% lower in the MNR females compared to the control *ad libitum* fed females (P=0.017; Fig. 2D). Nanoparticle gene therapy treatment did not change fetal weight of the restricted fetuses (P>0.05), but this result is unsurprising given the relative short time period between treatment and sample collection. Furthermore, there was no significant effect of fetal sex on fetal weight, nor on any of the other outcomes measured. Maternal-fetal interface weight was lower between MNR and control dams (P=0.031; Fig. 2E), and not changed by nanoparticle gene therapy treatment (P>0.05). Maternal-fetal interface efficiency (fetal to maternal-fetal interface weight ratio) was not different between either diet or nanoparticle gene therapy treatment (P>0.05; Fig. 2F).

**Fig. 2.**
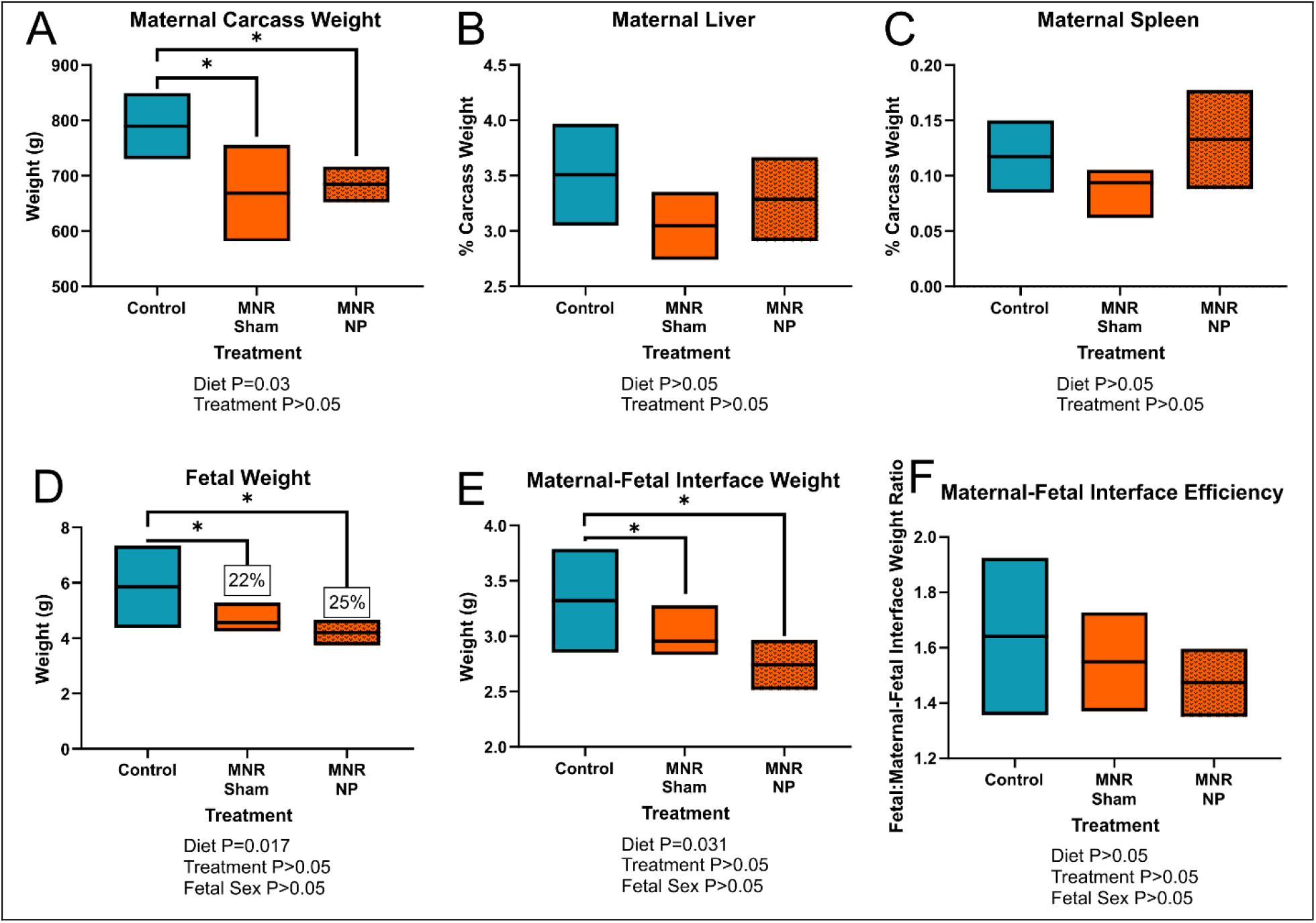
Effect of maternal nutrient restriction (MNR) diet and nanoparticle (NP) treatment on mid-pregnancy maternal, fetal and maternal-fetal interface weight. **A**. At time of sacrifice, maternal carcass weight (maternal weight minus total fetal and maternal-fetal interface weight) was lower in the MNR group compared to control diet group. **B**. Maternal liver weight as a percentage of carcass weight was not different with either diet or nanoparticle treatment. **C**. Maternal spleen weight at a percentage of carcass weight was not different with either diet or nanoparticle treatment. **D**. MNR resulted in approximately 22-25% reduction in fetal weight at mid-pregnancy, and there was no effect of placental nanoparticle gene therapy treatment in restricted fetuses. **E**. MNR reduced maternal-fetal interface weight at mid-pregnancy, which was also not affected by nanoparticle gene therapy treatment. **F**. There was no difference in maternal-fetal interface efficiency (fetal to maternal-fetal interface weight ratio) with either MNR diet or nanoparticle gene therapy treatment. Fetal sex was not a significant factor for any of the fetal/placental outcomes. *n* = 7 control dams (21 fetuses/placentas), 5 MNR-sham dams (14 fetuses/placentas) and 7 MNR-nanoparticle dams (19 fetuses/placentas). Data are estimated marginal means ± 95% confidence interval. P values calculated using generalized estimating equations with Bonferroni post hoc analysis. Asterix denote a significant difference of P≤0.05.

### Nanoparticle treatment had no effect on gross placental morphology but reduced placental labyrinth interhaemal distance in the maternal nutrient restricted group

Analysis of mid-sagittal cross-sections of the maternal-fetal interface showed no increased signs of necrosis or gross placental defects with neither MNR nor nanoparticle gene therapy treatment (Supplemental Fig. S7). There was also no difference in the sub-placenta/decidua area (P>0.05; Supplemental Fig. S7A), placenta area (P>0.05; Supplemental Fig. S7B), or total mid-sagittal area of the maternal-fetal interface (P>0.05; Supplemental Fig. S7C), with either diet or treatment. Double-label IHC was used to identify placental labyrinth trophoblast and fetal capillaries (Supplemental Fig. S1). In the restricted-sham placentas, trophoblast volume density was increased (P=0.01; Fig. 3A), but there was no difference in maternal blood space (MBS) volume density (Fig. 3B), or fetal capillary volume density (Fig. 3C), when compared to control. For the restricted nanoparticle gene therapy treated placentas, trophoblast volume density was not different when compared to control (Fig. 3A), but fetal capillary volume density was increased when compared to control and restricted sham (P=0.004; Fig. 3C). Most importantly, the interhaemal distance between maternal and fetal circulation was reduced in the restricted nanoparticle gene therapy treated placentas (P=0.043; Fig. 3D) indicating the possibility for greater oxygen diffusion.

**Fig. 3.**
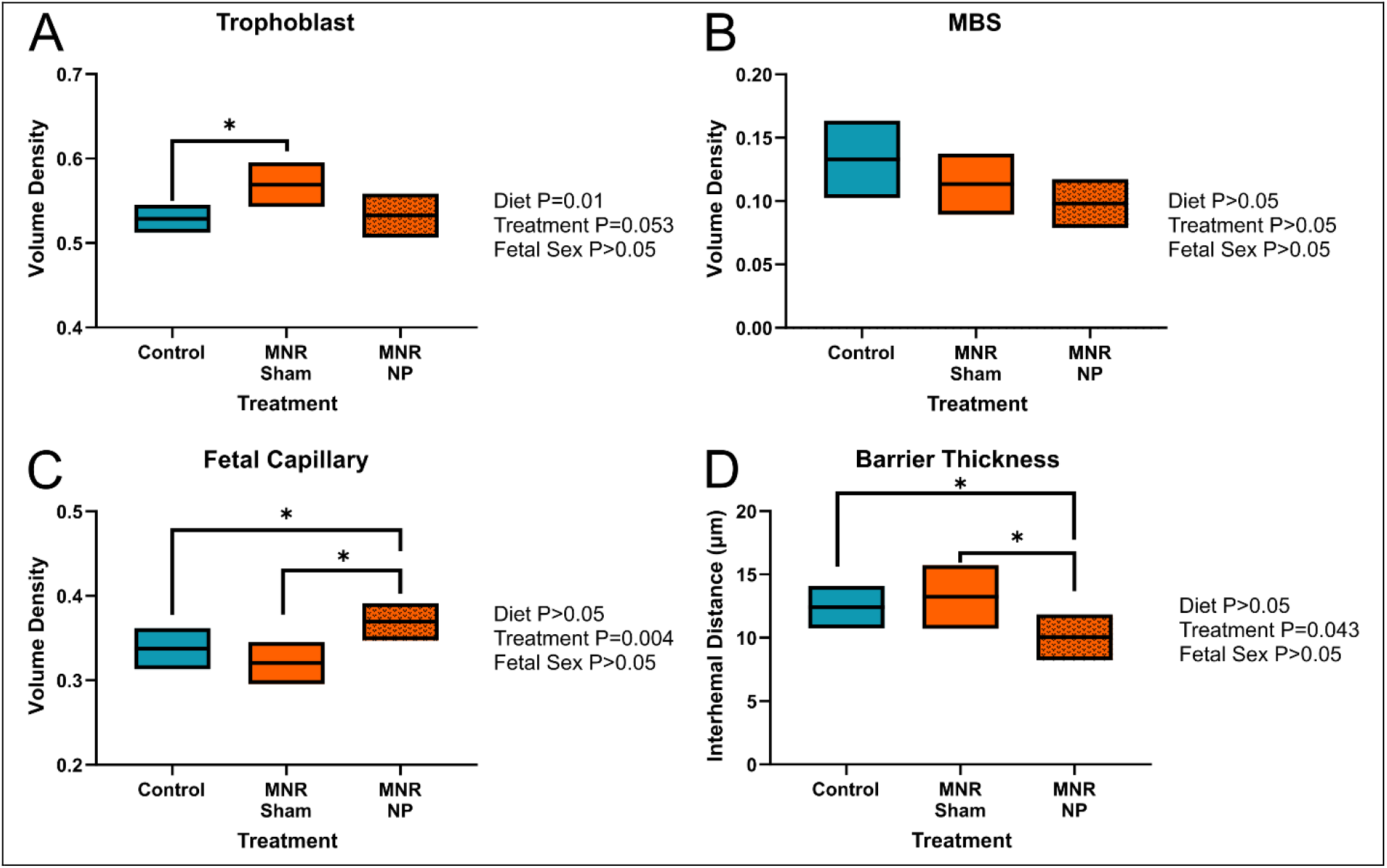
Effect of maternal nutrient restriction (MNR) diet and nanoparticle (NP) treatment on mid-pregnancy placenta microstructure. **A**. Compared to control, trophoblast volume density was increased in the restricted-sham placentas but not different in the restricted nanoparticle gene therapy treated placentas. **B**. There was no change to the maternal blood space (MBS) volume density with either diet or nanoparticle treatment. **C**. Nanoparticle gene therapy treatment of MNR placentas increased fetal capillary volume density compared to both control and restricted sham placentas. **D**. There was a reduction in the interhaemal distance (distance between maternal blood space and fetal circulation) in restricted nanoparticle gene therapy treated placentas compared to control and restricted sham placentas. There was no affect of fetal sex on any of the outcomes assessed. *n* = 7 control-sham dams (21 placentas), 5 MNR-sham dams (14 placentas) and 7 MNR-nanoparticle dams (19 placentas). Data are estimated marginal means ± 95% confidence interval. P values calculated using generalized estimating equations with Bonferroni post hoc analysis. Asterix denote a significant difference of P≤0.05.

### Placenta nanoparticle treatment increased fetal glucose concentrations and placenta expression of glucose and amino acid transporters

There was no difference in maternal glucose concentrations between control and MNR diet dams at time of post mortem (Fig. 4A). Despite no difference in fetal weight with nanoparticle gene therapy treatment, glucose concentrations were increased in the restricted nanoparticle gene therapy treated fetuses compared to control and restricted-sham fetuses (P=0.025; Fig. 4B). In restricted sham placentas, mRNA expression of glucose transporters *Slc2A1* was not different compared to control (Fig. 5A), however there was a reduction in expression levels between restricted nanoparticle gene therapy treated placentas and restricted sham (P=0.049; Fig. 5A). Western blot analysis of Slc2A1 protein expression showed no difference in global expression levels (Fig. 5B), however immunohistochemical staining for Slc2A1 showed relocalization of Slc2A1 to the apical membrane of the syncytium in the restricted nanoparticle gene therapy treated placentas (Fig. 5C). Expression of *Slc2A3* was reduced in restricted sham placentas compared to control (P=0.011; Fig. 5D), but increased in the restricted nanoparticle gene therapy treated placentas compared to restricted sham and comparable to control levels (P=0.002; Fig. 5D). There was also increased protein expression of Slc2A3 in the restricted nanoparticle gene therapy treated placentas compared to both control sham and restricted sham (P=0.005; Fig. 5D).

**Fig. 4.**
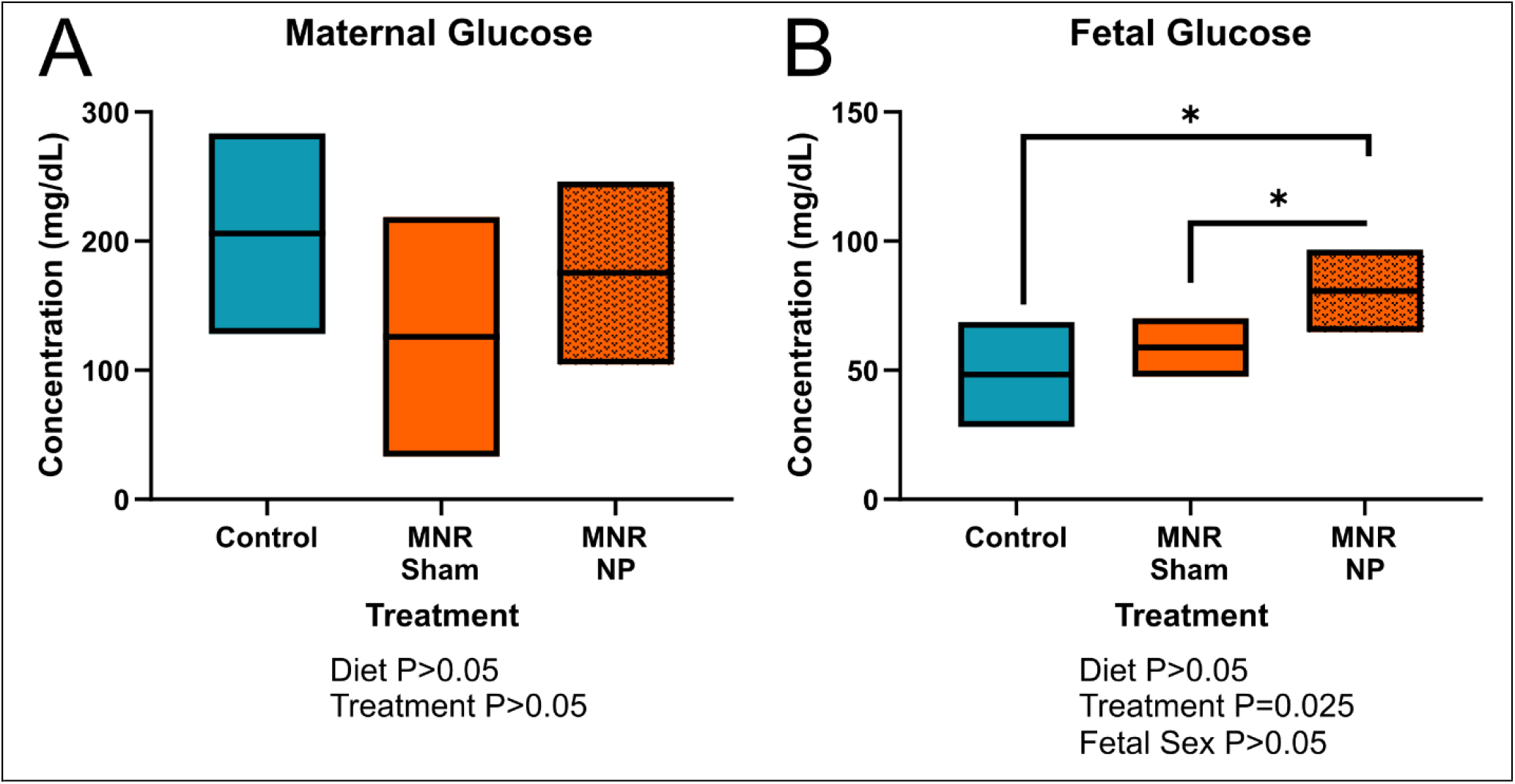
Effect of maternal nutrient restriction (MNR) diet and nanoparticle (NP) treatment on maternal and fetal blood glucose concentrations. **A**. There was no difference in maternal blood glucose concentrations at mid pregnancy with either diet or nanoparticle treatment. **B**. Nanoparticle gene therapy treatment increased glucose concentrations in restricted fetuses compared to control and restricted sham fetuses. *n* = 6 control dams (14 fetuses), 5 MNR-sham dams (10 fetuses) and 7 MNR-nanoparticle dams (17 fetuses). Data are estimated marginal means ± 95% confidence interval. P values calculated using generalized estimating equations with Bonferroni post hoc analysis. Asterix denote a significant difference of P≤0.05.

**Fig. 5.**
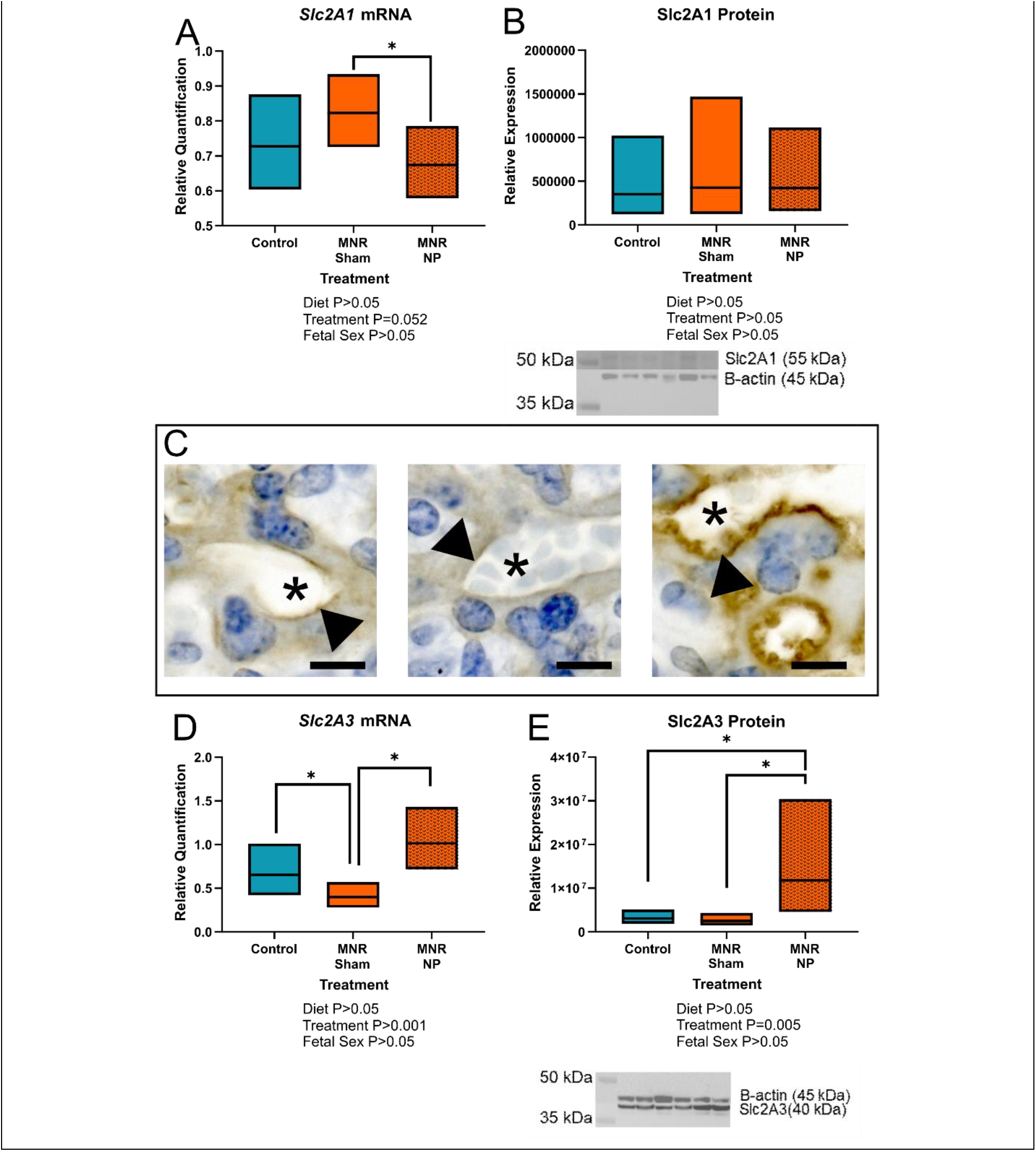
Effect of maternal nutrient restriction (MNR) diet and nanoparticle (NP) treatment on placental glucose transporter expression. **A**. Placenta expression of *Slc2A1* was reduced in MNR-nanoparticle gene therapy treated tissue compared to MNR-sham, but not different to control. **B**. There was no difference in protein expression of Slc2A1 in control-sham, MNR-sham or MNR-nanoparticle placentas. Representative western blot: lanes 1 & 2 = control-sham, lanes 3 & 4 = MNR-sham, lanes 5 & 6 = MNR-nanoparticle gene therapy treated. **C**. Representative images of immunohistochemical staining for Slc2A1 showing relocalization of Slc2A1 to the apical membrane of the placental syncytium (arrow head). Image 1 = control sham, image 2 = MNR-sham, image 3 = MNR-nanoparticle gene therapy treated. Asterix represents maternal blood space. **D**. Placenta expression of *Slc2A3* was reduced in restricted sham tissue compared to control, but increased back to normal levels in restricted nanoparticle gene therapy treated placentas. **E**. There was also increased protein expression of Slc2A3 in the restricted nanoparticle gene therapy treated placentas compared to control sham and restricted sham. Representative western blot: lanes 1 & 2 = control-sham, lanes 3 & 4 = MNR-sham, lanes 5 & 6 = MNR-nanoparticle gene therapy treated. *n* = 6 control dams (14 fetuses), 5 MNR-sham dams (10 fetuses) and 7 MNR-nanoparticle dams (17 fetuses). Data are estimated marginal means ± 95% confidence interval. P values calculated using generalized estimating equations with Bonferroni post hoc analysis. Asterix denote a significant difference of P≤0.05.

mRNA and protein expression of amino acid transporters Slc38A1 and Slc38A2 was also assessed in the placenta (Fig. 6). Placenta mRNA expression *Slc38A1* was not different between restricted sham and control placentas, but increased in restricted nanoparticle gene therapy treated placentas compared to control (P=0.016; Fig. 6A). For *Slc38A2*, expression was reduced in restricted sham placentas compared to control (P=0.048; Fig. 6B), but increased to normal levels in the restricted nanoparticle gene therapy treated placentas (P=0.011; Fig 6B.). Despite changes in gene expression, western blot analysis showed no changes to protein expression of either Slc38A1 (Fig. 6C) or Slc38A2 (Fig. 6D).

**Fig. 6.**
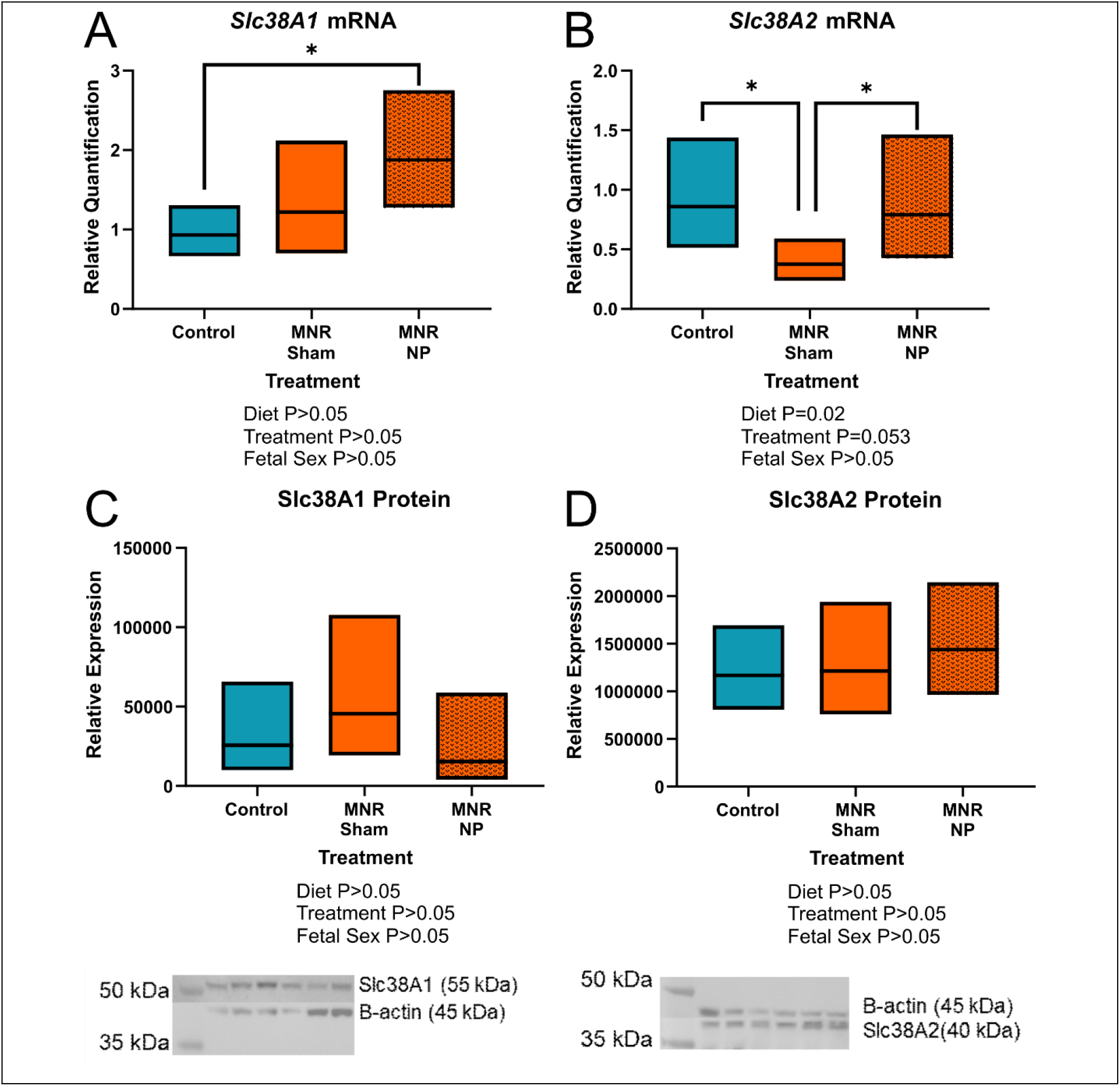
Effect of maternal nutrient restriction (MNR) diet and nanoparticle (NP) treatment placenta labyrinth amino acid transporter mRNA and protein expression. **A**. Placental expression of *Slc38A1* was increased in restricted nanoparticle gene therapy treated placentas compared to control. **B**. Placental expression of *Slc38A2* was reduced in MNR-sham placentas compared to control, but increased to normal levels in the MNR-nanoparticle gene therapy treated placentas. **C**. There was no difference in protein expression of Slc38A1 between the groups. Representative western blot: lanes 1 & 2 = control-sham, lanes 3 & 4 = MNR-sham, lanes 5 & 6 = MNR-nanoparticle gene therapy treated. **D**. There was no difference in protein expression of Slc38A2 between the groups. Representative western blot: lanes 1 & 2 = control-sham, lanes 3 & 4 = MNR-sham, lanes 5 & 6 = MNR-nanoparticle gene therapy treated. n = 6 control dams (14 placentas), 5 MNR-sham dams (10 placentas) and 7 MNR-nanoparticle dams (17 placentas). Data are estimated marginal means ± 95% confidence interval. P values calculated using generalized estimating equations with Bonferroni post hoc analysis. Asterix letters denote a significant difference of P≤0.05.

## Discussion

In the present study, we show efficient nanoparticle uptake in the mid-pregnancy guinea pig placenta and sustained transgene expression of *hIGF-1* in sub-placental trophoblast cells five days after nanoparticle gene therapy treatment. Nanoparticle gene therapy treatment was not toxic to pregnant dams as there was no difference in maternal weight or organ coefficients with treatment. Similarly, nanoparticle gene therapy treatment was not toxic to fetuses as no non-viable fetuses, that would have been lost due to treatment, were recorded at post mortems. Additionally, fetal weight was not lowered by treatment, and morphological assessment of the placenta did not show increased signs of necrosis or inflammation. Nanoparticle gene therapy treatment did however, increase fetal capillary volume density and reduce the interhaemal distance between maternal and fetal circulation in the growth restricted placenta, and increase fetal blood glucose concentrations by 37-66%. Additionally, there was changes in the expression of major glucose and amino acid transporters in the placenta. Such changes occurred within a short five-day time period, and indicates the potential for nanoparticle gene therapy treatment to have a positive effect on fetal growth with a longer duration of treatment.

The importance of IGF-1 in placental development and function is well established. Through binding with the IGF-1 Receptor, it signals to increase nutrient transport across the placenta as well as modulates placental vascular development [17]. In the current study, nanoparticle gene therapy treatment did not result in significant changes to fetal weight, nor gross placental morphology. However, given that the guinea pig pregnancy is 65-70 days, the lack of significant change to fetal weight is likely explained by the short time period between treatment and collection of samples. Nanoparticle gene therapy treatment was however, capable of increasing fetal capillary volume density. Angiogenesis is an essential component of placenta growth and development and IGF-1 is known to influence placental angiogenesis across gestation [35]. Furthermore, we have previously shown in the surgical ligated mouse model of FGR, increased *hIGF-1* expression increases the number of fetal vessels in the placenta [11]. Additionally, nanoparticle gene therapy reduced the placental interhaemal distance between the maternal blood space and fetal circulation. In human term placentas, IGF-1 treatment has been shown to inhibit the release of vasoconstrictors thromboxane B_2_ and prostaglandin F_2α_, indicating the ability for IGF-1 to enhance placental vasodilation and allow for increased blood supply to the placenta [36]. Thus, it is possible a similar mechanism is being activated in the current study, resulting in reduced interhaemal distance worth further investigation.

Glucose is an essential nutrient and energy source for the fetus, predominantly obtained from the mother [37]. Thus, adequate placental transfer of glucose from maternal circulation to fetal circulation is paramount to supporting appropriate fetal growth. In the present study, direct placental nanoparticle gene therapy treatment resulted in increased fetal glucose concentrations five days after administration indicating the ability to rapidly enhance placental glucose transport to the fetus within a short period of time. We have previously shown that increased *hIGF-1* expression in the placenta increased glucose transporter expression, either at the mRNA or protein level, in mice and human in vitro models [12, 20]. Furthermore, we have shown distinct re-localization of SLC2A1 to the apical membrane of the syncytiotrophoblasts in human placenta explants treated with IGF-1 nanoparticle gene therapy [12]; which has happened again with nanoparticle gene therapy treatment in the guinea pig placenta. Growth restricted nanoparticle gene therapy treated placentas also had increased expression of *Slc2A3* in both mRNA and protein. Human placental expression of SLC2A3 is highest in first trimester [38], and is thought to function as a rapid, high volume glucose transporter under conditions where extracellular glucose is lower [39], like in the first trimester when the placenta is absent of a fully formed maternal circulatory system. Altogether, these results provide a snapshot of how increased placental expression of *hIGF-1* may positively influence glucose transporter expression to enable rapid glucose uptake to the fetus.

In the placenta, amino acid transport is predominantly supported by the system A and L transporters [15]. System A transporters include Slc38A1 and Slc38A2, which are sodium-dependent [40]. In humans, reduced placental amino acid transport has been associated with down regulation of Slc38A2 expression in FGR [41]. The present study supports this finding, as there was a reduction in mRNA expression of Slc38A2 in the restricted-sham placentas compared to control. More importantly, nanoparticle gene therapy treatment of the restricted placentas returned expression levels of Slc38A2 to normal, and increased mRNA expression of Slc38A1. We have consistently shown that increasing placental expression of *hIGF-1* impacts the expression of Slc38A1 and Slc38A2 in the human placenta trophoblast cell line - BeWo cells, and surgically ligated mice placentas [19]. However, in these studies, there was also change to protein expression, not observed in our study of guinea pig placentas, but possibly again be due to the short time period in which nanoparticle treatment occurred. Never-the-less, our data indicates a positive effect on amino acid transporters with nanoparticle gene therapy likely to increase placental amino acid transport capacity under FGR conditions.

Development of an effective therapeutic to treat FGR during pregnancy is of high importance as FGR significantly contribute to the rates of stillbirth and miscarriage, and is also often associated with other fetal conditions, such as congenital heart defects [42]. Additionally, FGR is strongly associated with the developmental programming of adult diseases [43]. However, the implementation of treatments for FGR is complicated by the need to ensure the health and safety of both mother and offspring. In our current study, we found no toxic effects of short-term nanoparticle gene therapy treatment on either mother or fetuses; there was no difference in maternal or fetal weigh between nanoparticle and sham treated. Furthermore, our polymer-based nanoparticle gene therapy is designed to target the placenta which is discarded after birth, and represents the ideal site for an interventional therapy as there is no potentially long-term consequences to maternal health. Polymer-based systems have been shown to elicit less of an immune response and are safer than viral-based delivery systems [44-46], thus mitigating against concerns regarding immunogenicity and off-target affects [47]. Furthermore, a broad range of bioactive molecules including plasmids, siRNAs, mRNAs, proteins and oligonucleotides can be delivered with polymers [48-51] and as such, offers a promising platform for continuing to develop targeted strategies to treat the placenta and prevent FGR during pregnancy.

Our findings, that nanoparticle gene therapy treatment results in rapid changes to glucose transporter expression and increases fetal glucose concentrations, highlights the huge translational potential this treatment could have in human pregnancies. However, in the present study, gene therapy was injected directly to the placenta in order to maximize delivery and placental uptake. Clinically this method of delivery is sub-optimal and therefore, future studies utilizing intravenous gene therapy delivery and enhancing the placental homing capabilities are required. Overall, despite no significant changes to fetal weight or growth trajectories, short-term treatment of growth restricted placentas with the *hIGF-1* nanoparticle gene therapy resulted in increased fetal capillary volume density, reduced interhaemal distance and increased expression of glucose transporters associated with increased fetal glucose concentrations. Such outcomes indicate a potential positive effect on fetal growth worth investigating further with longer treatment.

## Contributions

RLW conceived the study, performed experiments, analyzed data and wrote manuscript. KKS and KL performed experiments, analyzed data and edited manuscript. MKG and CLD created polymer, analyzed data and edited manuscript. HNJ obtained funding, conceived the study and edited manuscript. All authors approve final version of manuscript.

## Ethics approval

Animal care and usage was approved by the Institutional Animal Care and Use Committees at Cincinnati Children’s Hospital and Medical Center (Protocol number 2017-0065) and University of Florida (Protocol number 202011236).

## Competing Interests

The authors have declared that no competing interest exists

## Data availability

All data needed to evaluate the conclusions in the paper are present in the paper and/or the Supplementary Materials.

## Funding

This study was funded by Eunice Kennedy Shriver National Institute of Child Health and Human Development (NICHD) award R01HD090657 (HNJ).

## Notes

### Competing Interest Statement

The authors have declared no competing interest.

### Summary of Updates

Have updated data to include changes in placental glucose and amino acids transporters in response to nutrient restriction and nanoparticle treatment Have removed data on effects to fetal liver gene expression

