## Supplemental Material for "Nanoparticle-mediated transgene expression of *insulin-like growth factor 1* in the growth restricted guinea pig placenta increases placenta nutrient transporter expression and fetal glucose concentrations"

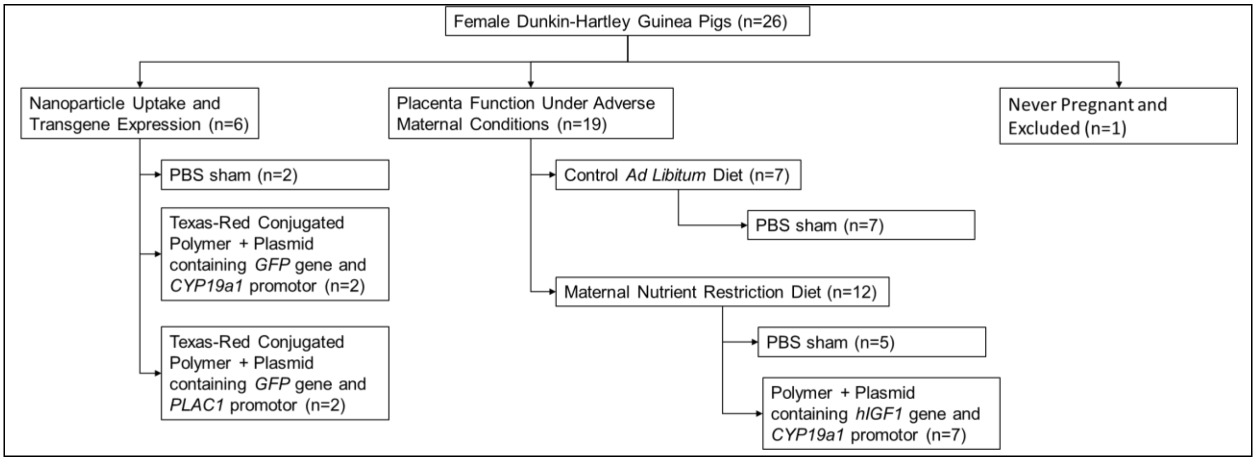


**Scheme S1** General schematic of experimental design and animal allocations

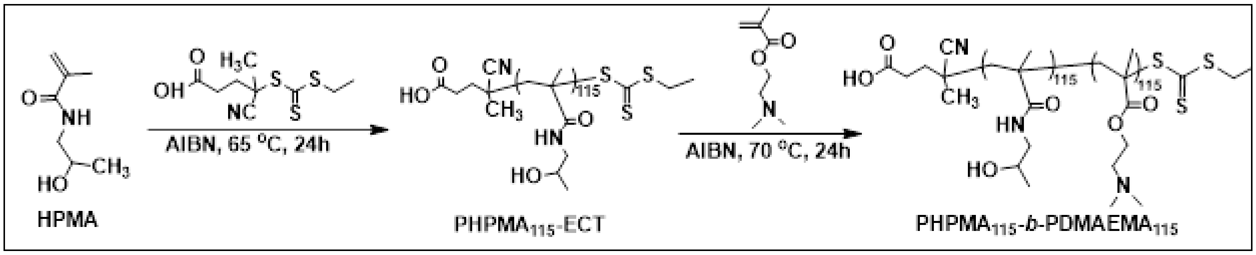


**Scheme S2** General schematic of the synthesis of a diblock copolymer of $PHPMA_{115}$-b-$PDMAEMA_{115}$ by RAFT polymerization.

**Supplemental Table S1. Primer sequences for qPCR**

| **Target** | **Forward Primer** | **Reverse Primer** |
|---|---|---|
| *Slc2A1* | CCTGCAGTTTGGCTACAACA | GTCTGGTTGTAGAACTCCTCG |
| *Slc2A3* | GATCCCACAAATACCAACAG | CAGAAACCACAGTGAAGATG |
| *Slc38A1* | GCTAGGAGAACAAGTCTTTG | CATACCAGGCTGAAAATGTC |
| *Slc38A2* | CAGAACATTGGAGCTATGTC | CAGATACCACAATCCAGTTG |
| *β-actin* | CGCGAGAAGATGACCCAG | TAGCACAGCCTGGATAGCAA |
| *Rsp20* | GTGAAAGGACCCGTTCGCAT | CTTCACCACAGGGCGTTTTC |

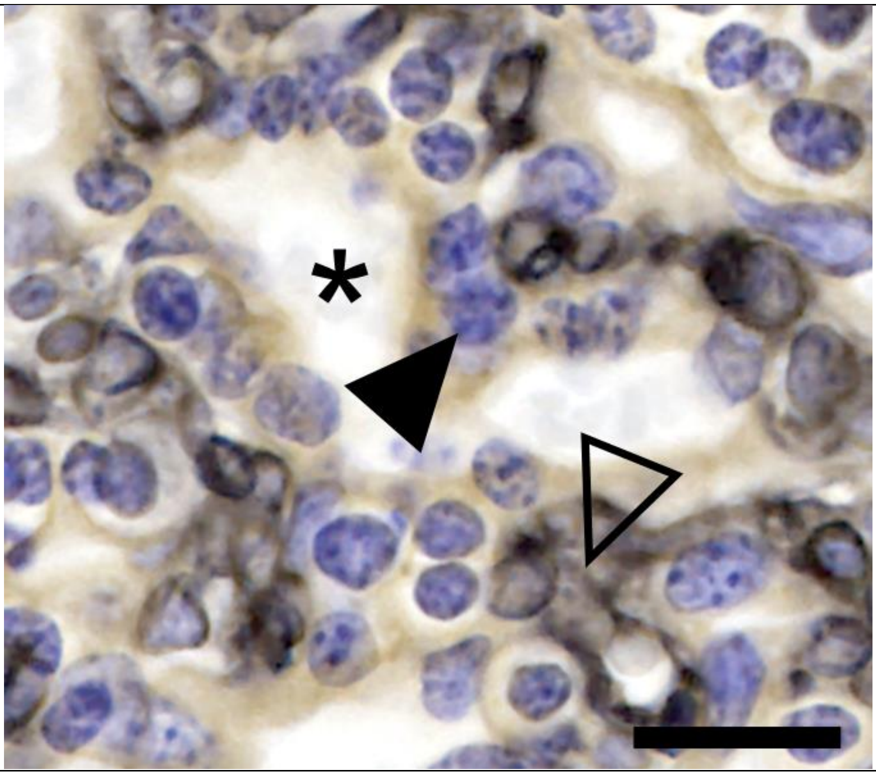

**Supplemental Figure S1.** Representative image of double-label immunohistochemistry in the guinea pig placental labyrinth. Fetal capillary endothelium was labelled with an anti-vimentin antibody (black: open arrow). Placental trophoblast was labelled with an anti-cytokeratin antibody (brown: closed arrow). Asterix indicates maternal blood space. Scale bar = 20 µm.

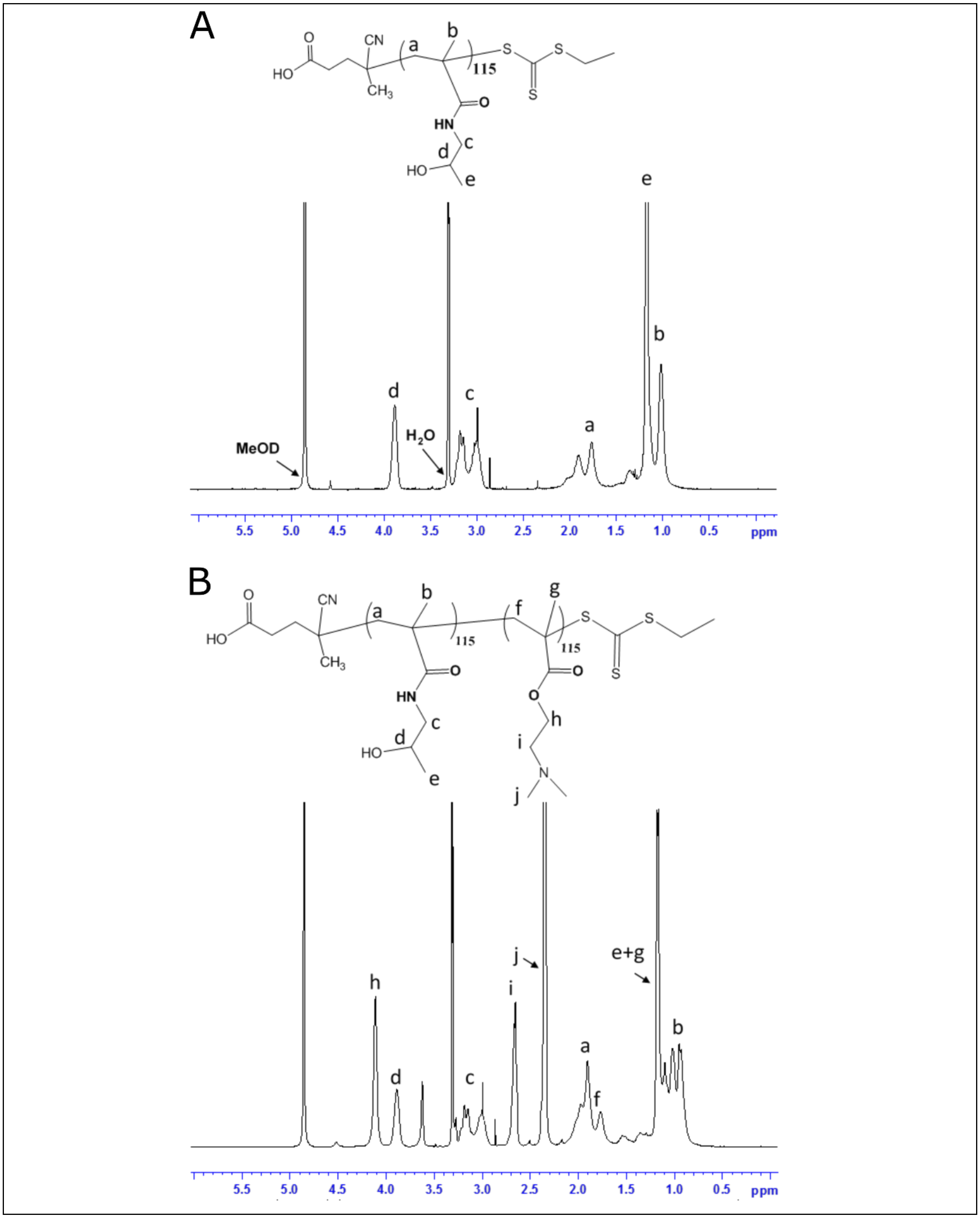


**Supplemental Figure S2.** Confirmation of successful deblock polymer formation. 1H NMR spectra of PHPMA115-ECT in Methanol-d4 (**A**). The peaks from RAFT macro-CTA are not assigned because they are relative insignificant and also merge with PHPMA protons. 1H NMR spectra of PHPMA115-b-PDMAEMA115 in Methanol-d4 (**B**).

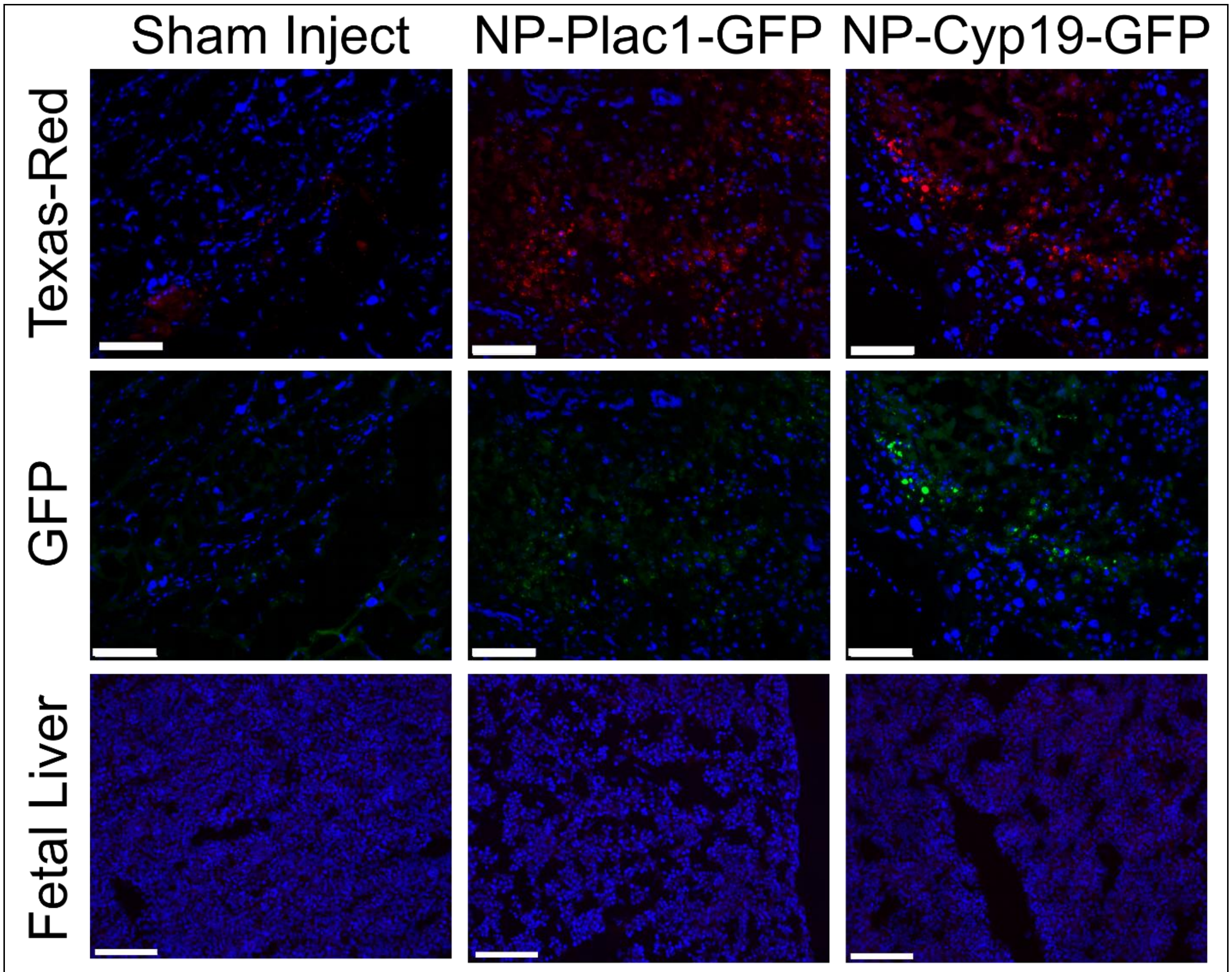


**Supplemental Fig. S3. Fluorescent microscopy of nanoparticle (NP-Plac1-GFP and NP-Cyp19-GFP) localization and transgene expression in the guinea pig placenta and fetal liver.** Conjugation of the HPMA-DMEAMA copolymer with a Texas Red fluorophore (Red) confirmed nanoparticle uptake in the guinea pig placenta 30 h after intra-placental injection. Analysis of green fluorescent protein (GFP; Green) fluorescence showed only the Cyp19, and not the Plac1, promotor was recognized by guinea pig trophoblast cells. Fluorescence of neither Texas Red nor GFP was found in fetal liver tissue confirming the inability for nanoparticle to cross the placenta and enter fetal circulation. Nuclei counter-stained with Dapi (Blue). Scale bar = 0.1 mm.

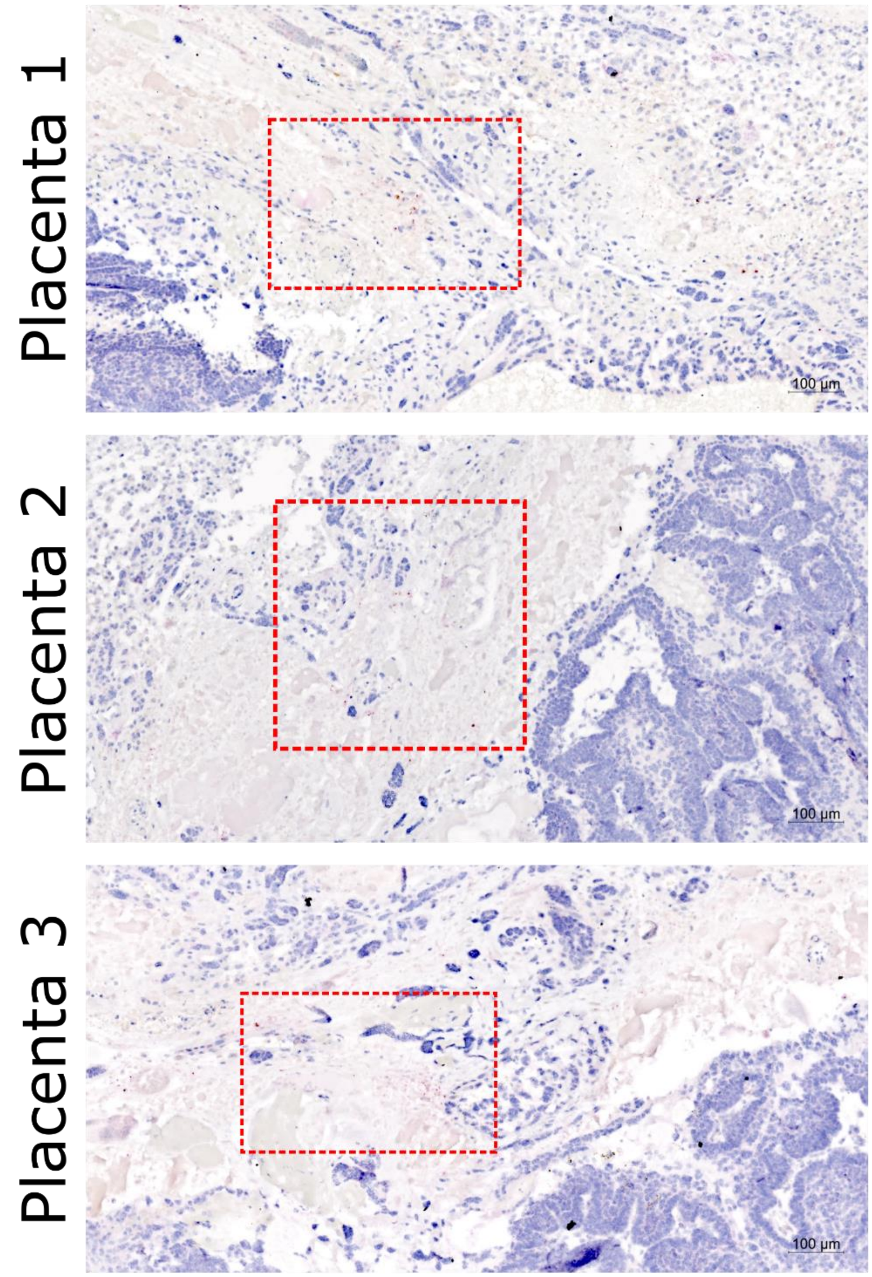


**Supplemental Figure S4.** In situ hybridization (ISH) for plasmid-specific mRNA expression in the guinea pig placenta five days after intra-placental nanoparticle treatment. Despite only one placenta receiving direct placental injection, ISH detected plasmid-specific mRNA in all placentas from the litter including placentas located on the opposite uterine horn.

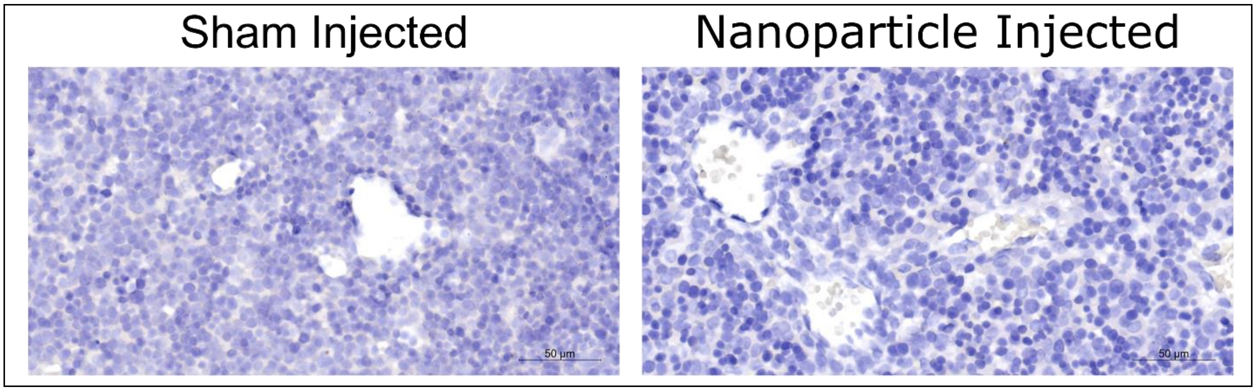


**Supplemental Figure S5.** In situ hybridization (ISH) for plasmid-specific mRNA expression in guinea pig fetal liver five days after intra-placental nanoparticle treatment. ISH did not detect the presence of plasmid-specific mRNA in the fetal liver further confirming the inability for nanoparticle to cross the placenta and enter fetal circulation.

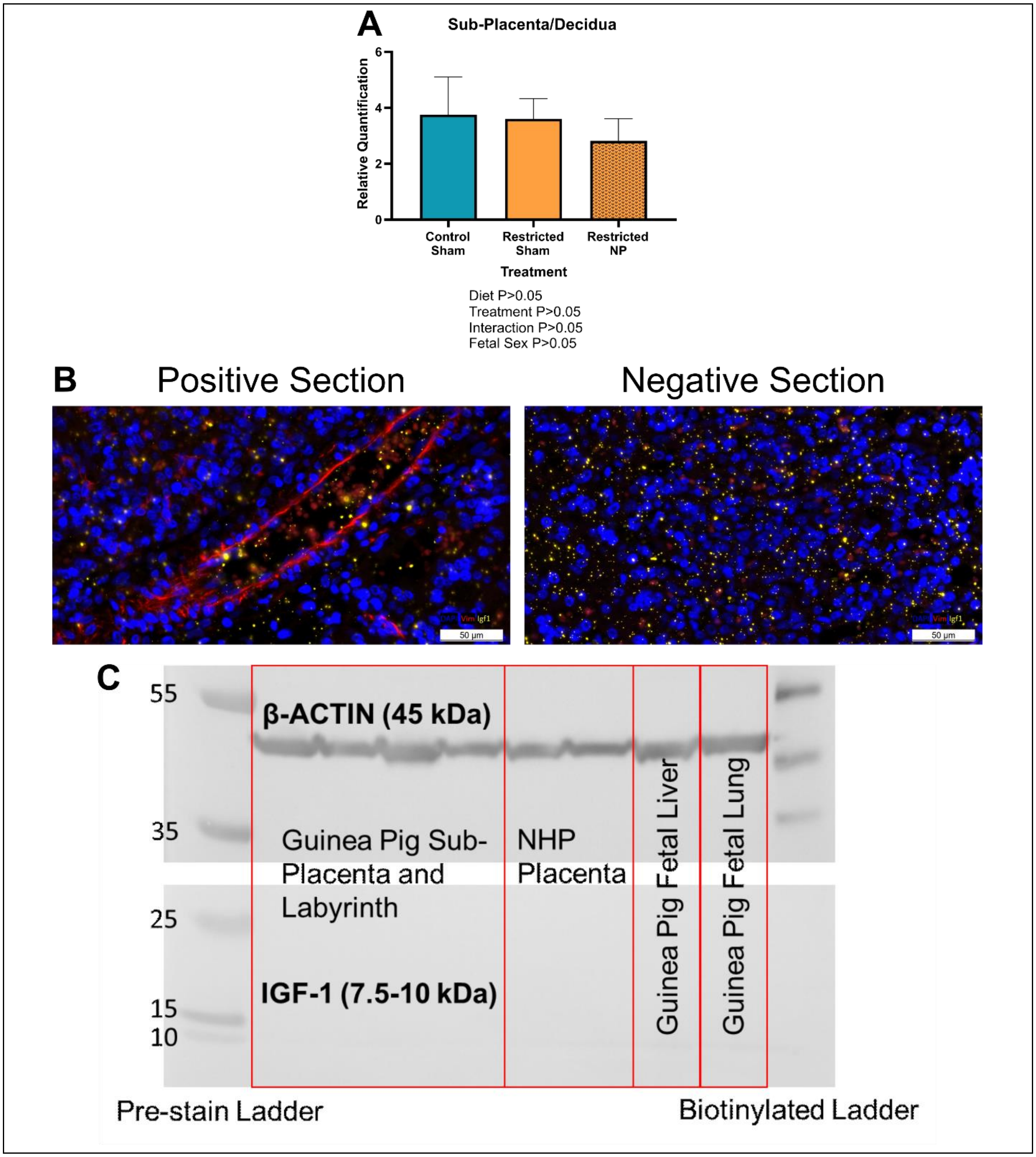


**Supplemental Figure S6.** Analysis of insulin like 1 growth factor (Igf-1) in guinea pig placenta by qPCR, immunohistochemistry and western blot. qPCR analysis of *Igf-1* in guinea pig sub-placenta/decidua tissue was not affected by diet or treatment (**A**); there was no expression (amplification at CT>35) in the guinea pig placental labyrinth. Double-label immunohistochemistry for Vimentin (red; positive antibody to confirming correct staining procedure) and Igf-1 (yellow) showed no positive staining for Igf-1 (**B**). Similarly, there was no positive detection of Igf-1 at the expected molecular weight using western blot (**C**). β-actin was used as a positive control to confirm correct blotting procedure.

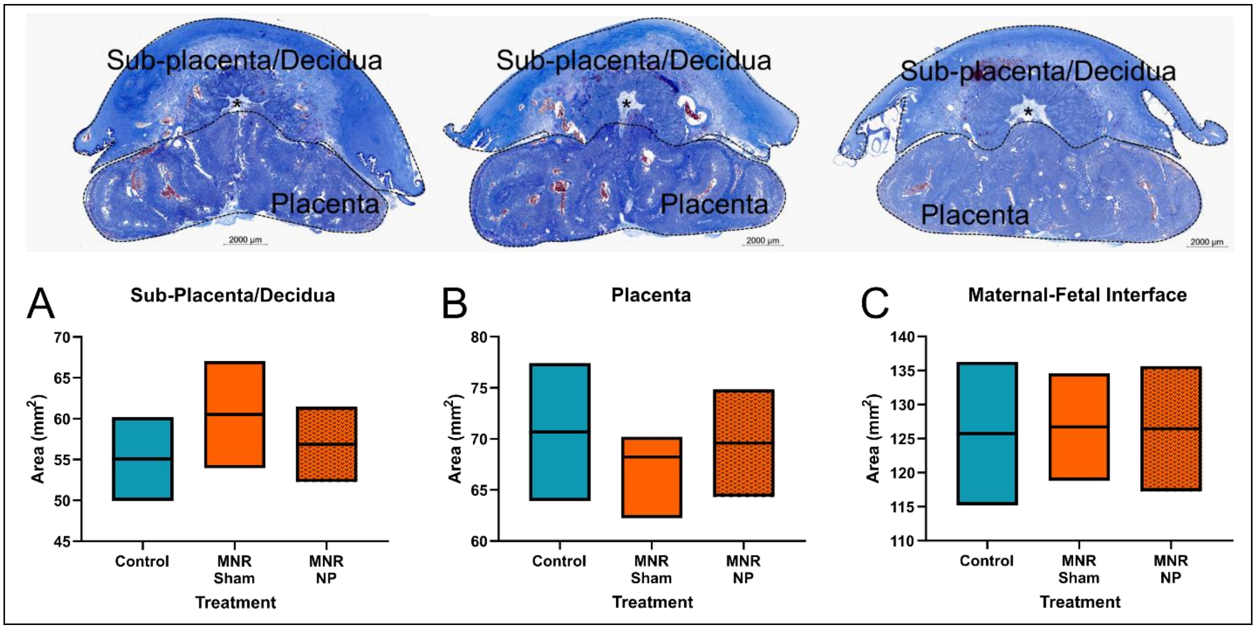


**Supplemental Fig. S7. Effect of maternal nutrient restriction (MNR) diet and nanoparticle (NP) treatment on mid-pregnancy gross placental morphology.** There were no indicators of increased necrosis or inflammation in the placenta with intra-placental treatment. Neither MNR diet nor nanoparticle treatment effected sub-placenta/decidua area (**A)**, labyrinth area (**B)**, nor labyrinth weight (**C**). Images are representative mid-sagittal cross sections of a control-sham placenta (left), MNR-sham placenta (middle) and MNR nanoparticle treated placenta (right). *n* = 7 control-sham females (21 placentas), 5 MNR-sham females (14 placentas) and 7 MNR-nanoparticle females (19 placentas). Data are estimated marginal means ± 95% confidence interval. P values calculated using generalized estimating equations with Bonferroni post hoc analysis.
